# Nonsteroidal anti-inflammatory drugs alter the microbiota and exacerbate *Clostridium difficile* colitis while dysregulating the inflammatory response

**DOI:** 10.1101/391607

**Authors:** Damian Maseda, Joseph P. Zackular, Bruno Trindade, Leslie Kirk, Leslie J. Crofford, Patrick D. Schloss, Jennifer Lising Roxas, V.K. Viswanathan, Gayatri Vedantam, Lisa M. Rogers, Mary K. Washington, Eric P. Skaar, David M. Aronoff

## Abstract

*Clostridium difficile* infection (CDI) is a major public health threat worldwide. The use of nonsteroidal anti-inflammatory drugs (NSAIDs) is associated with enhanced susceptibility to and severity of nosocomial CDI; however, the mechanisms driving this phenomenon have not been elucidated. NSAIDs alter prostaglandin (PG) metabolism by inhibiting cyclooxygenase (COX) enzymes. Here, we found that treatment with the NSAID indomethacin prior to infection altered the microbiota and dramatically increased mortality and intestinal pathology associated with CDI in mice. We demonstrate that in *C. difficile-infected* animals, indomethacin lead to PG deregulation, an altered proinflammatory transcriptional and protein profile, and perturbed epithelial cell junctions. These effects were paralleled by an increased recruitment of intestinal neutrophils and CD4^+^ cells. Together, these data implicate NSAIDs in perturbation of the gut microbiota and disruption of protective COX-mediated PG production during CDI, resulting in altered epithelial integrity and associated immune responses.

---

*Clostridium difficile* is the most commonly reported nosocomial pathogen in the United States and an urgent public health threat worldwide(1). *C. difficile* infection (CDI) manifests as a spectrum of gastrointestinal disorders ranging from mild-diarrhea to toxic megacolon and/or death, particularly in older adults(2). The primary risk factor for CDI is antibiotic treatment, which perturbs the resident gut microbiota and abolishes colonization resistance(3). However, factors other than antibiotic exposure increase the risk for CDI and cases not associated with the use of antimicrobials have been on the rise(4). Defining mechanisms whereby non-antibiotic factors impact CDI pathogenesis promises to reveal actionable targets for preventing or treating this infection.

Recently, several previously unappreciated immune system, host, microbiota, and dietary factors have emerged as modulators of CDI severity and risk. The food additive trehalose, for example, was recently shown to increase *C. difficile* virulence in mice and the widespread adoption of trehalose in food products was implicated in the emergence of hypervirulent strains of *C. difficile*(5). Similarly, excess dietary zinc has a profound impact on severity of *C. difficile* disease in mice, and high levels of zinc alter the gut microbiota and increase susceptibility to CDI(6). Importantly, there has been a growing body of evidence for the essential role of the innate immune response and inflammation in both protection against and pathology of CDI(7–9). Mounting a proper and robust inflammatory response is critical for successful clearance of *C. difficile*, and the immune response can be a key predictor of prognosis(3, 10). In this context, specific immune mediators can facilitate both protective and pathogenic responses through molecules like IL-23 and IL-22, and an excessive and dysregulated immune response is believed to be one of the main factors behind post-infection complications.

Epidemiological data have established an association between the use of nonsteroidal anti-inflammatory drugs (NSAIDs) and CDI(11). Muñoz-Miralles and colleagues demonstrated that the NSAID indomethacin significantly increased the severity of CDI in antibiotic-treated mice when the NSAID was applied following inoculation and throughout the infection (Muñoz-Miralles *et al.*, in press, Future Microbiology, 2018), and indomethacin exposure is associated with alterations in the structure of the intestinal microbiota(12, 13). NSAIDs are among the most highly prescribed and most widely consumed drugs in the United States(14), particularly among older adults(15) and have been implicated in causing spontaneous colitis in humans(16, 17). They act by inhibiting cyclooxygenase (COX) enzymatic activity, which prevents the generation of prostaglandins (PGs) and alters the outcome of subsequent inflammatory events. Prostaglandins, and among those especially PGE_2_, are important lipid mediators that are highly abundant at sites of inflammation and infection, and support gastrointestinal homeostasis and epithelial cell health (18). NSAID use has been associated with shifts in the gut microbiota, both in rodents and humans(19–22), but these shifts have not been explored in the context of CDI.

In this report, we deployed a mouse model of antibiotic-associated CDI to examine the impact of exposure to indomethacin prior to infection with *C. difficile* on disease severity, immune response, intestinal epithelial integrity, and the gut microbiota. These investigations revealed that even a brief exposure to an NSAID prior to *C. difficile* inoculation dramatically increases CDI severity, reduces survival, and increases pathological evidence of disease. Inhibition of PG biosynthesis by indomethacin altered the cytokine response and immune cell recruitment following CDI, enhancing intestinal tissue histopathology and allowing a partial systemic bacterial dissemination by dismantling intestinal epithelial tight junctions. Additionally, indomethacin treatment alone significantly perturbed the structure of the gut microbiota. These findings support epidemiological data linking NSAID use and CDI, and caution against the overuse of NSAIDs in patients at high risk for *C. difficile*, such as older adults.

## Results

### Indomethacin worsens C. difficile infection in mice and increases mortality

To determine the extent to which pre-exposure to NSAIDs influences the natural course of CDI, mice were treated with indomethacin for two days prior to inoculation with *C. difficile* (Fig. 1A). We infected C57BL/6 female mice with 1×10^4^ spores of the *C. difficile* NAP1/BI/027 strain M7404 following 5 days of pre-treatment with the broad-spectrum antibiotic, cefoperazone (Fig. 1A). This brief indomethacin treatment prior to CDI dramatically decreased cecum size, increased mortality rate from 20% to 80% (Fig. 1C) but did not significantly impact weight loss (Fig. 1D). Mice pre-treated with indomethacin and infected with *C. difficile* also displayed more severe histopathological evidence of cecal tissue damage compared to mice infected with *C. difficile* that were not exposed to the drug (Fig. 1E). Indomethacin-exposed and infected mice exhibited no change in the burden of *C. difficile* in the cecum (Fig. 1F), but their livers harbored significantly greater loads of mixed aerobic and anaerobic bacteria (Fig. 1G), suggesting that indomethacin pre-treatment compromised intestinal barrier function during CDI and fostered microbiota translocation to the liver.

**Fig. 1:**
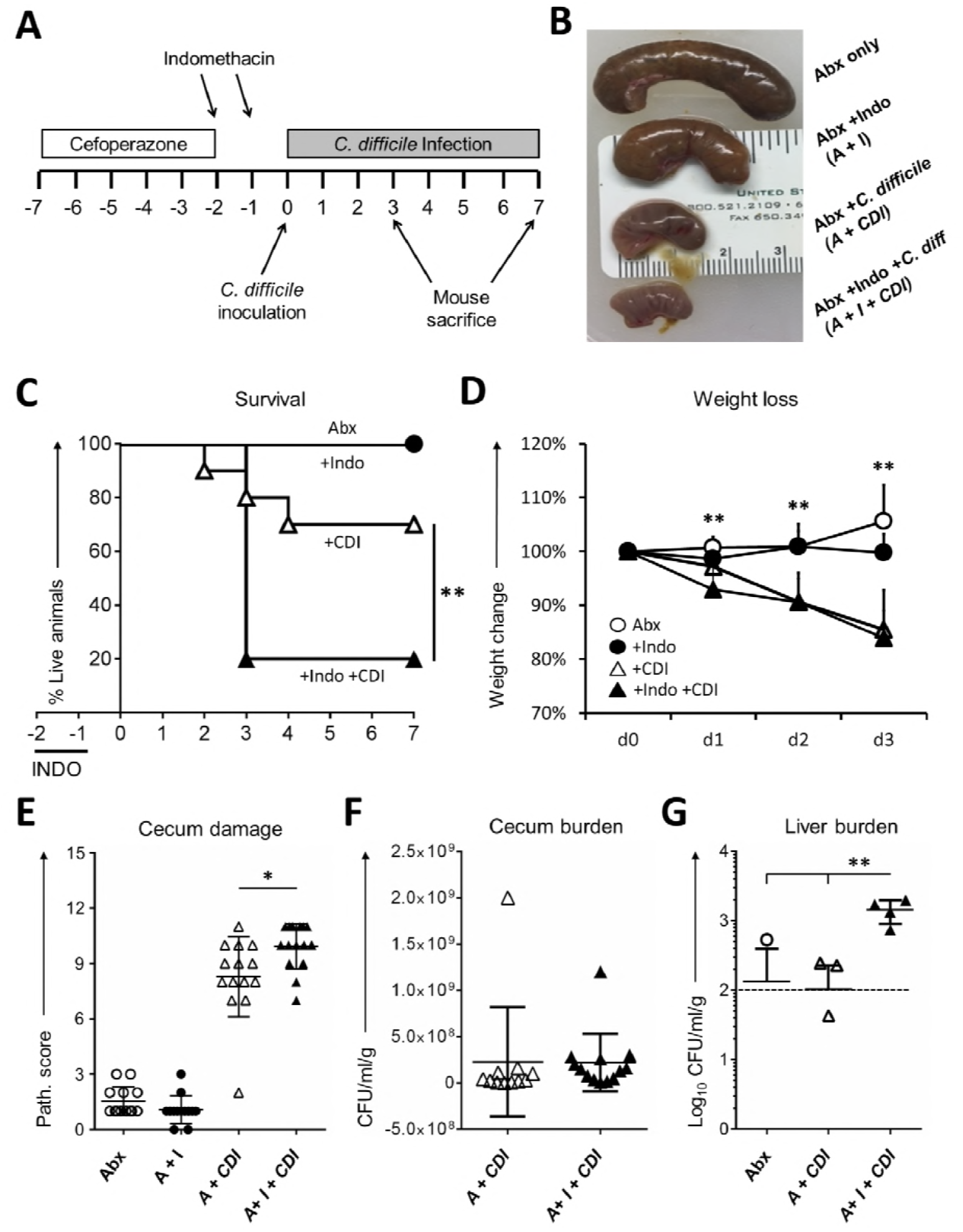
Indomethacin worsens *C. difficile* in mice. C57BL/6 mice were treated with cefoperazone for 5 days followed by 2 days of recovery and then challenged by gavage with 1×10^4^ spores of the NAP1 strain M7404. Animals received 2 doses of 10 mg/kg of indomethacin by gavage daily as indicated by the top arrows in **(A)**. Representative picture illustrating the macroscopic effects of the different treatments in the cecum **(B)**. Mice were monitored for survival (**C**), weight loss (**D**), and histopathologic severity of colitis (**E**). *C. difficile* bacterial burden was evaluated in the ceca of 12 mice/group **(F)**, and total aerobic + anaerobic bacterial burden in the liver of 5 mice/group **(G)** also at day 3 after infection, with the discontinuous line indicating limit of detection. **P<0.01 by Log-rank (Mantel-Cox) test for survival and *P<0.05, **P<0.01 by unpaired t test.

### Indomethacin alters the proportion of neutrophils and CD4^+^ T cells in mucosal-associated tissues during C. difficile infection

The mucosal immune response is an important factor in the clearance of and the pathology associated with CDI(10, 23–26). NSAIDs can disturb immune homeostasis within the gastrointestinal mucosa(27) and have been used to trigger immune-mediated colitis in mice(28). We determined the extent to which indomethacin altered immune cell populations in and around the gastrointestinal tract during CDI. Mice were euthanized by day 3 after infection and cells from the peritoneal cavity, mesenteric lymph nodes (mLN) and colonic lamina propria (cLP) were processed for flow cytometry analysis. CDI provoked an increase of neutrophil and CD4^+^ T cell numbers across all three compartments (Fig. 2). Focusing on the differences caused by indomethacin exposure prior to CDI, we found that neutrophils were significantly increased in the peritoneal cavity compared to CDI alone. This was paralleled by a similar overall trend in the mLN and colonic lamina propria (Fig. 2B). On the other hand, CD4^+^ T cells were slightly decreased in the mLN, but their numbers increased in the cLP (Fig. 2C), potentially due to a selective migration and/or proliferation in inflamed sites. Considering that IL-17 has been implicated in driving the neutrophilic inflammatory response to CDI (Nakagawa et al., 2016) and that Th17 cells and ILC3 cells are major sources of IL-17 during inflammatory responses, we evaluated the combined impact of indomethacin and CDI on these populations. Interestingly, larger numbers of CD4^+^RORγt+ (Th17) cells were found in the cLP, but not in the mLN (Fig. 2D). CDI also induced an expansion of ILC type 3 cell numbers in the cLP, but without significant alterations due to indomethacin pre-treatment (Fig. 2E). These data demonstrate that indomethacin pre-treatment exacerbates neutrophilic and Th17-type immune responses to CDI in the mouse.

**Fig. 2:**
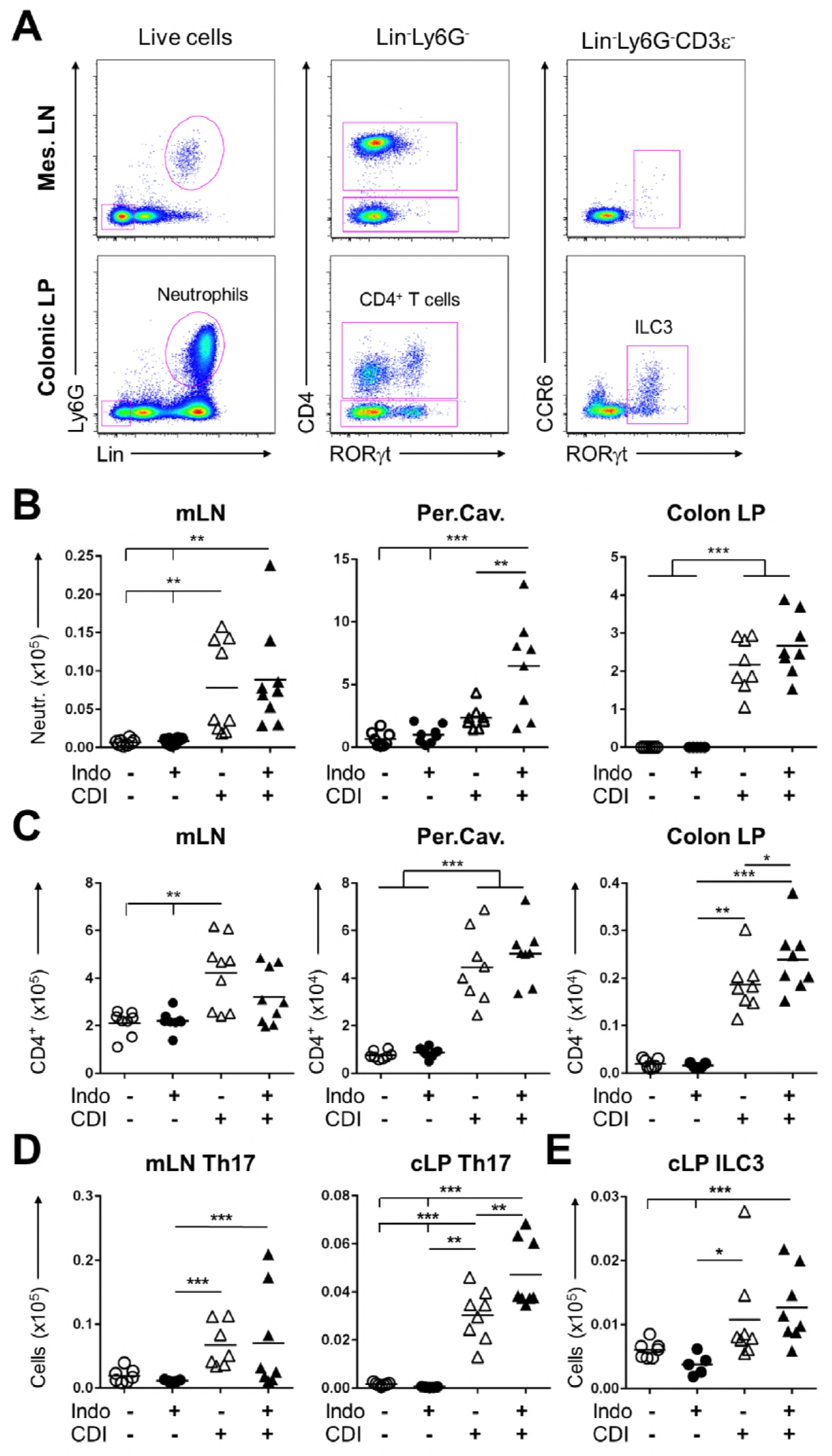
Indomethacin alters the proportions of neutrophils and CD4^+^ T cells in mucosal associated tissues during CDI. Mice were treated as previously described and were euthanized 3 days after infection. The colon lamina propria (cLP), mesenteric lymph nodes (mLN) and peritoneal cavity (Per.Cav.) were collected for analysis by flow cytometry (n = 8-10/group). **(A)** Representative flow plots from d3 CDI mice infected but not treated with indomethacin depicting the gating used to identify neutrophils (Lin^+^Ly6G^+^), CD4^+^ T cells (Lin^−^CD4^+^) and ILC group3 cells (Lin^−^Ly6G^−^CD4^−^RORγt^+^) in different organs. **(B)** Neutrophils and **(C)** CD4^+^ T cells numbers (x10^6^). Quantification of the analysis of mLN and cLP CD4^+^RORγt^+^ Th17 cells **(D)** and ILC type 3 (Lin^−^CD3ε^−^RORγt^+^) cells **(E)**. The middle line is presented as average. One-way ANOVA with Turkey’s correction was used to evaluate significant differences among all groups: *P<0.05, **P<0.01, ***P<0.001.

### Indomethacin dysregulates the expression of genes involved in prostaglandin metabolism and inflammatory peptides during CDI

CDI induces extensive transcriptional changes in the intestines that generally result in protective responses that restrain bacterial spread and mitigate induced intestinal epithelial pathology(3, 29). To examine the impact of indomethacin on this response, we interrogated transcriptional changes related to inflammatory responses in the cecum following indomethacin pre-treatment followed by CDI. We noted significant alterations, both positive and negative, in the inflammatory gene transcriptome of the cecum in mice infected with *C. difficile* following brief indomethacin exposure compared with *C. difficile*-inoculated mice that were not treated with the NSAID (Fig. 3B-3D). Notably, indomethacin pre-treatment followed by CDI significantly upregulated several genes involved in innate immune cell activation and recruitment like *Il1b, Cxcl3, Csf3, Cxcl1*, while it downregulated *Cd4, Tlr5* and *Tgfb2* (Fig. 3.C and D).

**Fig. 3:**
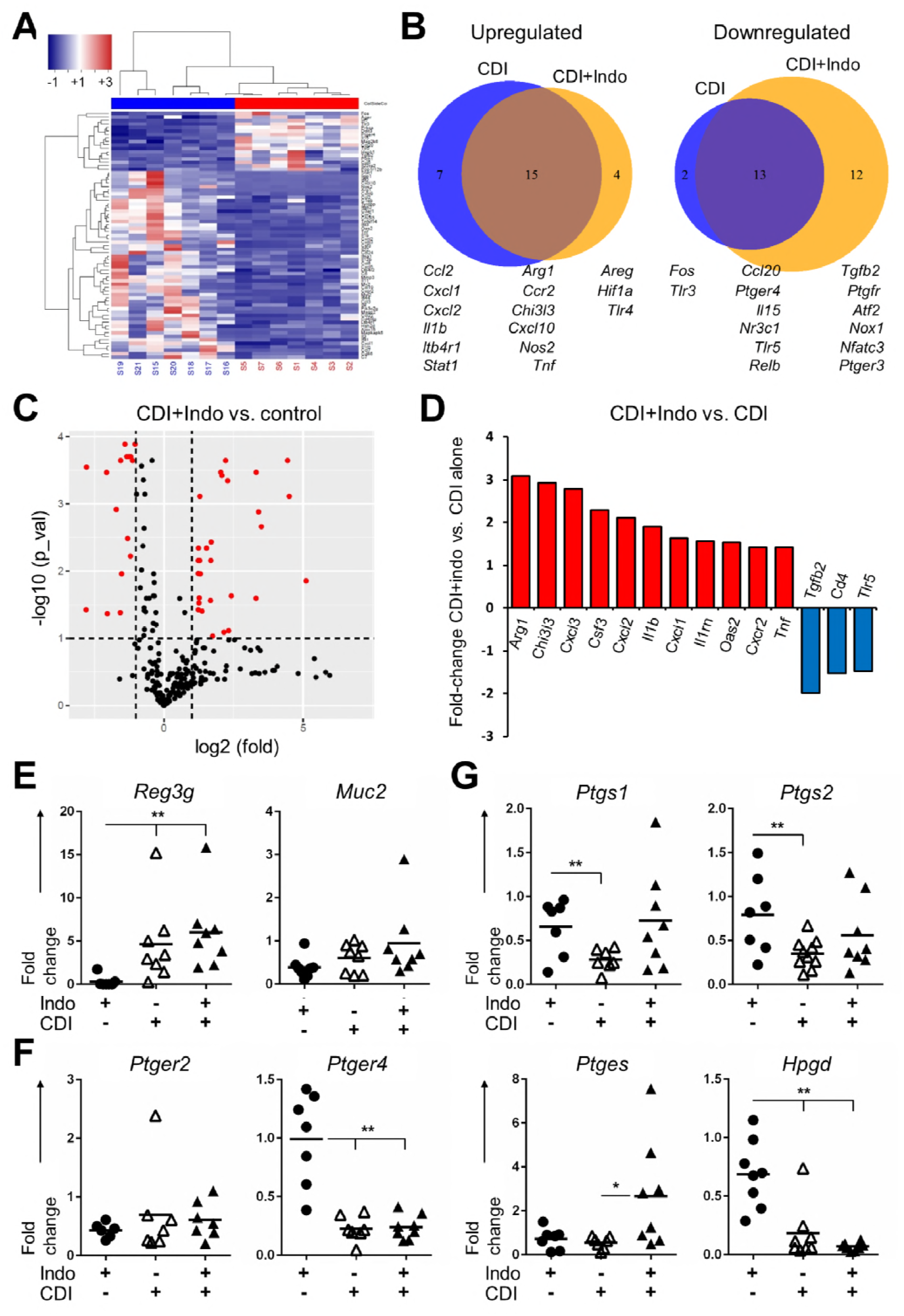
Prostaglandin inhibition by indomethacin inhibits an intestinal protective PGE_2_-mediated response to *Clostridium difficile* and induces damage driven by innate immune cells. **(A)** Representative clustering showing relative mRNA expression comparing the groups that received CDI alone versus control (cefoperazone only). **(B)** Venn diagram depicting overlap in gene up- and down-regulation upon CDI or CDI+indomethacin pre-treatment compared to control mice. Size of circles is proportional to number of genes. **(C)** Volcano plot of CDI+indomethacin versus control, n=12 samples/group. Red dots in the volcano plots are significantly differently expressed genes that are either under- or overexpressed. **(D)** Summary of genes that depict the highest up- (red) and downregulation (blue) fold differences when comparing the CDI+indomethacin versus CDI alone groups. Ceca of mice belonging to mice undergoing treatment were used to obtain mRNA, generate cDNA and perform RT-PCR (n = 8/group) at day 3 post-CDI. **(E)** Relative mRNA expression of intestinal markers of inflammation and protection *Reg3g and Muc2* **(F)** PGE_2_ receptors EP2 and EP4 (*ptger2 and ptger4*), and **(G)** enzymes controlling PGE_2_ metabolism COX1, COX2, mPGES and 15-PGDH (*Ptgs1, Ptgs2, Ptges* and *Hpgd*). See also Fig. S1. *P<0.05 and **P<0.01 in a 1-way ANOVA.

To further characterize the impact of NSAIDs on the immune response during CDI, we explored the impact of indomethacin on intrinsic mechanisms of host defense in the gastrointestinal tract. Specifically, we focused on the Gram-positive selective antimicrobial peptide, REG3γ, and mucin; two host intestinal defense factors that have been shown to be important for the control of gastrointestinal infections(30). We confirmed by qRT-PCR that CDI upregulated *Reg3g* transcription, while *Muc2* transcript levels were not significantly altered following indomethacin treatment (Fig.3E). To evaluate if PGE_2_ synthesis and signaling were altered due to infection or indomethacin treatment, we analyzed the expression of genes encoding PGE_2_ receptors and the enzymes involved in PGE_2_ metabolism. The transcription of the PGE_2_ receptor gene *Ptger4* was severely suppressed upon CDI, but indomethacin did not significantly exacerbate this suppression (Fig. 3F). Infection C. difficile suppressed colonic expression of the COX-1 and COX-2 encoding genes *Ptgs1* and *Ptgs2*, respectively (Fig. 3G). Notably, indomethacin pre-treatment prevented this down-modulation and simultaneously induced the expression of the *Ptges* gene, which encodes a major synthase for PGE_2_ (Fig. 3G). What is more, indomethacin further reduced expression of the PGE_2_ inactivating enzyme 15-hydroxyprostaglandin dehydrogenase (*Hpgd* gene; Fig. 3G). This selective inhibition of *Ptgs1* and *Ptgs2* transcription, together with inhibition of *Hpgd* and enhanced *Ptges* are consistent with the paradoxical *increase* in PGE_2_ concentrations observed 72 hours following infection (Fig. S1). Together, these data demonstrate that indomethacin pre-treatment increases innate immune cell activation and recruitment, while also leading to PG dysregulation.

### Indomethacin increases intestinal inflammation by upregulating a combined myeloid-recruitment and response in the cecum

Following the observation that indomethacin pre-treatment significantly altered cellular and transcriptional immune responses during CDI, we sought to determine the impact of this drug on tissue-level inflammatory protein expression during infection. Infected mice (either exposed to indomethacin or not) were euthanized and ceca were harvested at day 3 post-CDI. Whole tissue homogenates were used to measure the concentration of a panel of inflammation-related proteins and were normalized to total protein content per cecum (Fig. 4). The protein levels observed largely supported the transcriptomics results from our previous results, confirming what has already been reported for CDI regarding IL-1 β and immune mononuclear cell recruitment and activation proteins like CCL3, CXCL2 and CCL4. Interestingly, IL-6-class cytokines (IL-6, LIF) were among the most enhanced by indomethacin pre-treatment, consistent with what has been found in humans infected with *C. difficile*(24, 31, 32). Together with the increase in IL-1 β, and consistent with the above results showing enhanced Th17 responses, these data implicate an exacerbated IL-17A-related response caused by indomethacin. In contrast, some type-1-associated inflammatory molecules like IL-12p40 were downregulated by indomethacin pre-treatment.

**Fig. 4:**
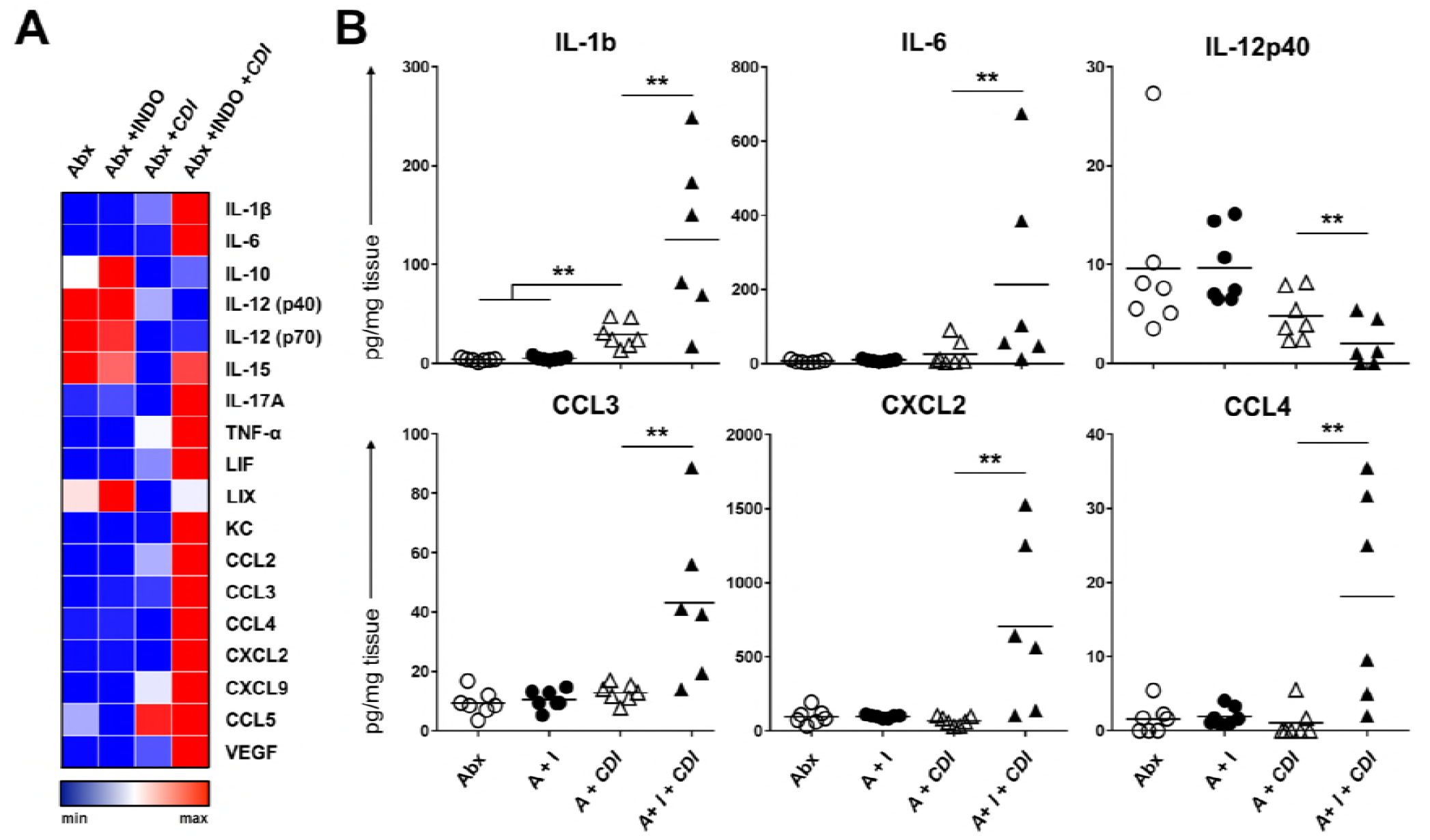
Indomethacin treatment enhances the inflammatory milieu in ceca of mice infected with *C. difficile*. Mice were treated as in figures 3-4 and their ceca collected at day 3 after infection. **(A)** Protein expression levels in homogenates from individual ceca were measured by Luminex and plotted on a log-2 scale. N =7-8/group. All values are provided in pg protein/cecum protein content **(B)**. Selected pro-inflammatory cytokines and myeloid cell-recruiting chemokines plotted to depict range of variation.

### Indomethacin perturbs colonic epithelial cell junctions of C. difficile-infected mice

The observations describing the increased bacterial translocation (Fig. 1G), together with the increased local PGE_2_ levels (Fig. 1H) and inflammatory molecules (Fig. 3), lead us to investigate whether the integrity of the intestinal epithelial barrier was compromised due to indomethacin pre-treatment during CDI. We assessed the impact of indomethacin on the integrity of colonic epithelial junctions of *C. difficile*-infected mice via transmission electron microscopy and immunofluorescence staining of tight junction (TJ) and TJ-associated proteins. Intestinal epithelial cells (IECs) of uninfected, cefoperazone-treated and uninfected, cefoperazone and indomethacin pre-treated mice had uniform microvilli and intact cell junctions similar to mock-treated mice (Fig. 5A). *C. difficile* infection resulted in microvilli effacement of intestinal epithelial cells, but did not appear to cause gross structural alteration of the cell junctions. In contrast, indomethacin pre-treatment of *C. difficile-infected* mice triggered striking intestinal epithelial cell separation at the region of the TJs.

**Fig. 5:**
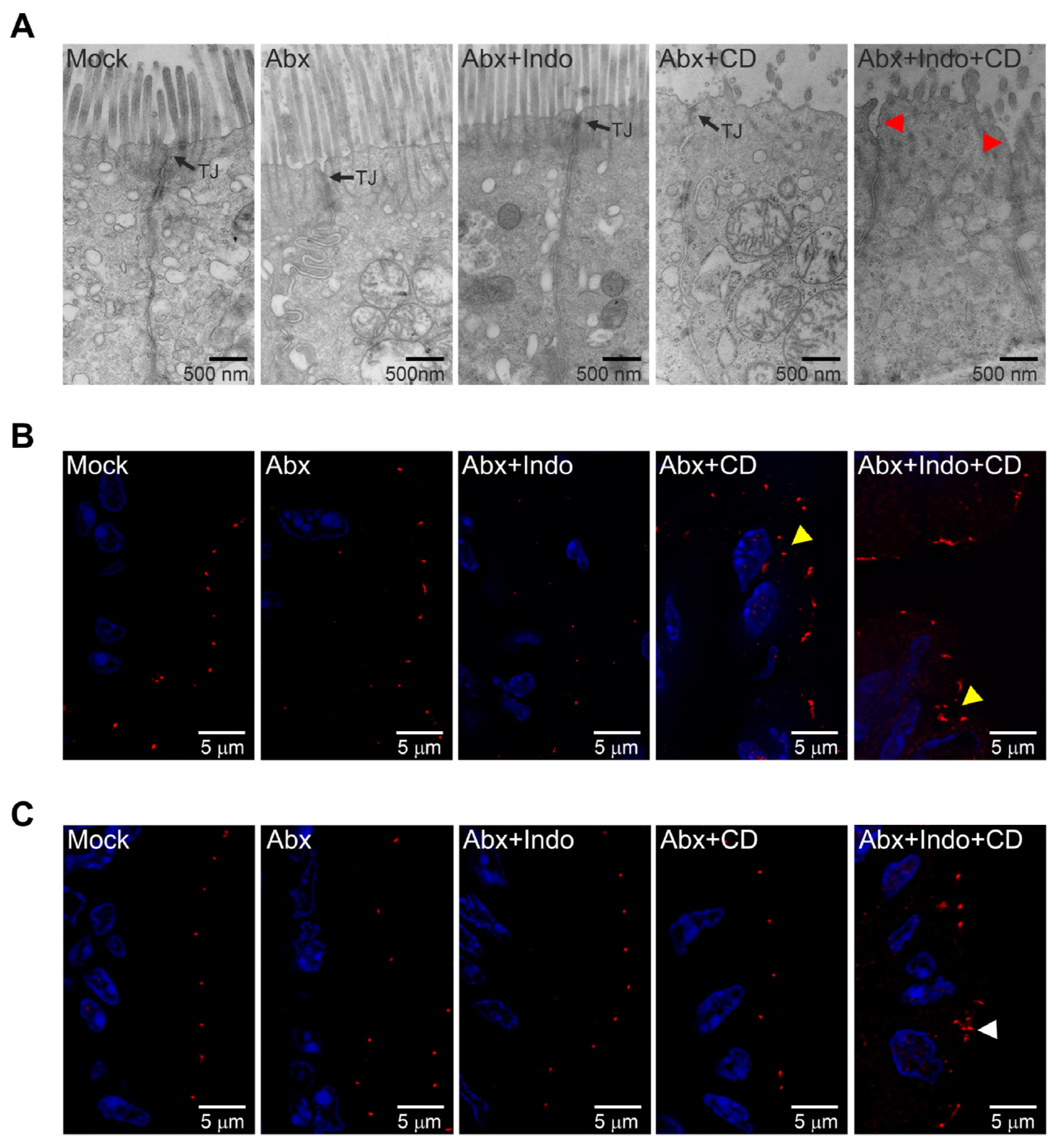
Indomethacin promotes relocalization of TJ-associated protein ZO1 and perturbs colonic epithelial cell junctions of *C. difficile-infected* mice. **(A)** Transmission electron micrographs showing lateral views of colonic mucosa from untreated control mice (Mock) or mice treated with the antibiotics cefoperazone alone (Abx), cefoperazone and indomethacin (Abx+Indo), cefoperazone and *C. difficile* (Abx+CD), or cefoperazone, indomethacin and *C. difficile* (Abx+Indo+CD). Arrows point to intact tight junctions (TJ). Red arrowheads point to TJ unzipping or separation. Mouse colonic tissues from the same groups above were stained for TJ-protein occludin **(B)** and TJ-associated protein ZO1 **(C)**. Occludin and ZO1 stain are pseudo-colored in red. DAPI (blue) was used to stain DNA. Yellow or white arrow heads indicate cytoplasmic relocalization of occludin and ZO1, respectively.

TJ complexes containing membrane-anchored occludin, claudins and junctional adhesion molecules (JAMS) attach to the perijunctional actomyosin ring via adaptor proteins such as zona occludens 1 (ZO1). Consistent with the intact cell junctions observed in IECs of uninfected mice, Occludin and ZO1 localized at the apex of lateral cell junctions (Fig. 5B and 5C). In contrast, CDI resulted in occludin relocalization to the cytoplasm of epithelial cells. ZO1 redistribution to the cytoplasm, however, was observed only in *C. difficile-infected* mice that were previously treated with indomethacin. Collectively, our data suggest that indomethacin acts synergistically with *C. difficile* to alter the localization of occludin and ZO1 and perturb TJ integrity of intestinal epithelial cells *in vivo*.

### Indomethacin alters the intestinal microbiota composition without further reducing microbial community diversity after antibiotic treatment

The composition of the gut microbiota has a profound impact on the manifestation and clearance of CDI, as well as the virulence of *C. difficile* and the outcome of disease(33, 34). There is also evidence suggesting prominent off-target effects of pharmaceutical agents, such as NSAIDs, on the gut microbiota and gastrointestinal health (20, 35). To examine the impact of indomethacin on the murine gut microbiota, mice were treated with a two-day course of indomethacin and the microbial community was subsequently surveyed using 16S rRNA gene sequencing. One day post-treatment, mice given indomethacin showed no significant alteration in α-diversity (Fig. S2) but exhibited a significant shift in community structure compared to untreated mice (P<0.001; AMOVA) (Fig. 6A). To characterize differentially abundant taxa in indomethacin treated mice, we utilized the biomarker discovery algorithm LEfSe (linear discriminant analysis (LDA) effect size). Indomethacin treatment was associated with an enrichment in operational taxonomic units (OTUs) affiliated with the *Bacteroides* (OTU 1), *Akkermansia* (OTU 4), and *Parasutterella* (OTU 17) genera, and the *Porphyromonadaceae* (OTU 14) family (Fig. 6B-C). Moreover, we observed a significant decrease in OTUs affiliated with the *Turicibacter* (OTU 18) genus and *Porphyromonadaceae* (OTU 5) family following indomethacin treatment (Fig. 6B-C). To examine the longitudinal impact of indomethacin on the murine gut microbiota, we collected samples periodically for 11 days following administration of indomethacin. We observed significant differences in community structure up to 2 days following administration of indomethacin treatment and a significant enrichment of *Bacteroides* (OTU 1) could be detected as far as 11-days following treatment with indomethacin (Fig. 6C). Next, to determine how indomethacin may impact the microbiota in the context of antibiotic treatment, mice were again exposed to indomethacin for 2 days following 5 days of cefoperazone (0.5 mg/ml) treatment. Although cefoperazone treatment dramatically reduced overall community α-diversity in all mice, we detected significant alterations in community structure associated with co-treatment with indomethacin and cefoperazone that could be observed 11-days post-treatment (Fig. 6D; Fig. S3). Initial differences in microbial community structure were driven by a significant bloom in *Paenibacillus* (OTU 11), while *Akkermansia* (OTU 6) was significantly enriched in mice treated with indomethacin and cefoperazone on day 11 post-treatment (Fig. 6.E-F). Together these data suggest that indomethacin has a marked effect on the structure of the gut microbiota and these off-target effects likely contribute to disease exacerbation during CDI.

**Fig. 6:**
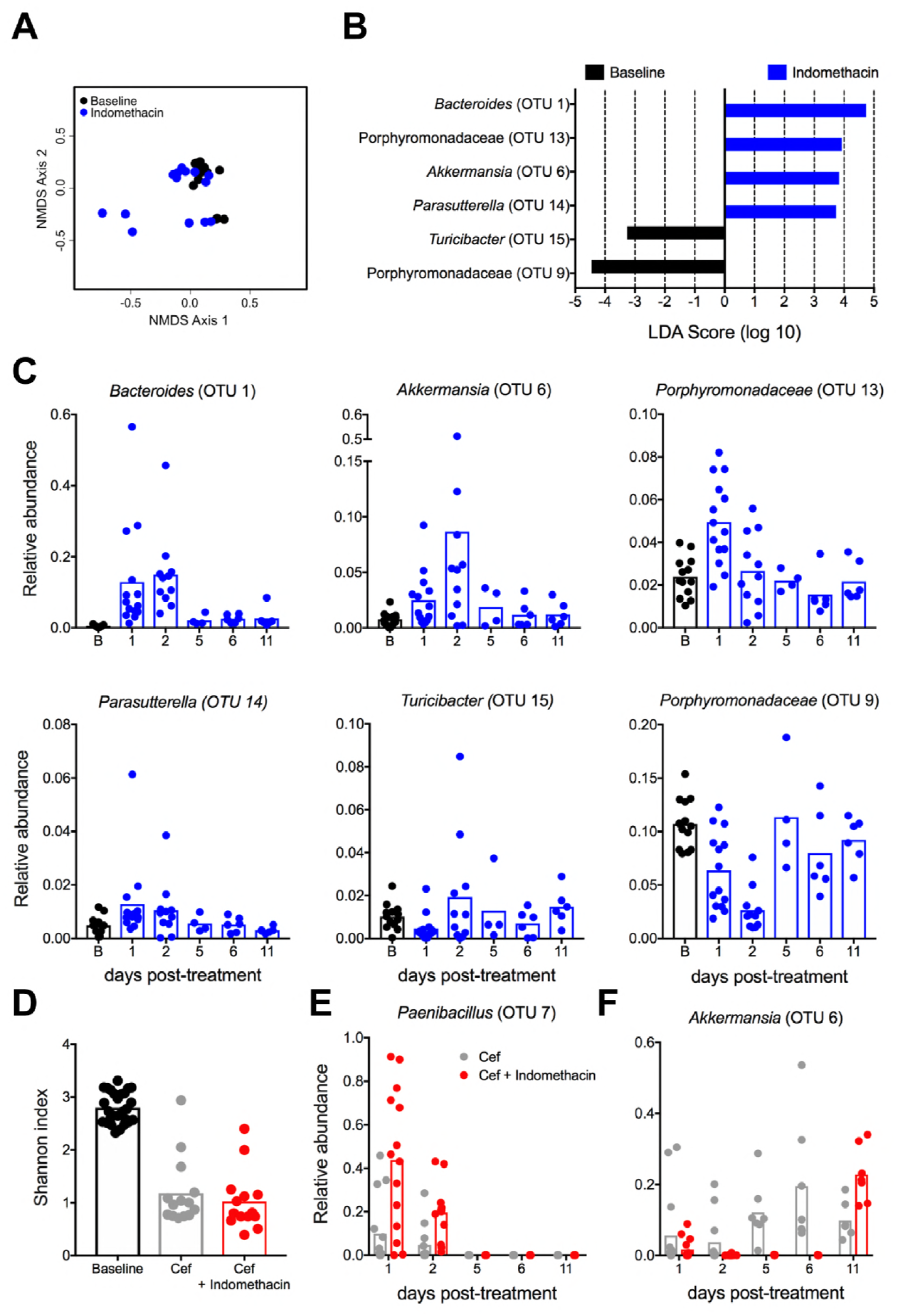
Indomethacin treatment alters the gut microbiota. (**A**) Non-metric multidimensional scaling (NMDS) ordination showing β-diversity as measured by Yue and Clayton’s measure of dissimilarity (Θyc) on day 1 post-indomethacin treatment. Significance between baseline (black) and indomethacin-treated (blue) samples measured using analysis of molecular variance (AMOVA) (P<0.001). (**B**) Differentially abundant taxa in baseline and indomethacin treated animals ranked by effect size. (**C**) Dynamics of recovery of differentially abundant taxa over an 11-day time course. B represents baseline microbiota pretreatment. Sample size (N): B = 13, d1 = 14, d2 = 11, d5 = 4, d6 = 6, d11 = 6. (**D**) Shannon diversity index for untreated (baseline; black), cefoperazone treated (grey), or cefoperazone and indomethacin treated (red) mice. See also Fig. S2. Dynamics of (**E**) *Paenibacillus* and (**F**) *Akkermansia* relative abundance following cefoperazone (grey) and cefoperazone + indomethacin treatment (red). See also Fig. S3.

## Discussion

CDI is the most commonly diagnosed cause of antibiotic-associated diarrhea and has surpassed methicillin-resistant *Staphylococcus aureus* as the most common healthcare associated infection in many US hospitals(36). Nearly 30,000 people die each year in the US from CDI (37). A major challenge of CDI is recurrence, which can impact 20-30% of patients and is associated with an increased risk of death(4, 38). One of the most promising treatments for CDI is fecal microbiota transplantation (FMT), which is estimated to be >80% effective in most studies(39, 40). However, problems with standardization, availability, and putative risks from FMT have made this form of therapy suboptimal(41). There continues to be a demand for effective approaches to limit CDI severity and to understand complications arising from the synergy of CDI with intestinal immune responses and drugs used to limit damaging inflammatory effects.

The NSAIDs are among the most commonly prescribed drugs in the US with more than 98 million prescriptions filled annually(14), and an estimated 29 million Americans using over-the-counter NSAIDs per year(42). As they prevent synthesis of endogenous PGs, NSAIDs can adversely affect intestinal health. Epidemiological studies reveal an association between CDI risk and the use of NSAIDs, underscored by a recent meta-analysis(11). The plausibility of a link between NSAID use and CDI is bolstered by the association between NSAID use and flare-ups of inflammatory bowel disease and the occasional occurrence of NSAID-induced colitis(43–46). Recent mouse studies have established that concomitant NSAID use exacerbates active CDI (Muñoz-Miralles *et al.*, in press, Future Microbiology, 2018).

Animal and human studies suggest that CDI induces local and systemic increases in PGs such as PGE_2_(47). Prostaglandin E_2_ is one of the most common and well-characterized PGs, which has long been known to have major effects on gastrointestinal health(48–51). COX-1-dependent production of PGE_2_ is gastroprotective, explaining why chronic NSAID use is associated with stomach ulcers, and why such ulcers can be prevented by administering the FDA-approved oral PGE analogue misoprostol to NSAID-treated patients(52). In addition, endogenous PGE_2_ production prevents gut epithelial cell death and promotes colonic tumor growth by directly inducing tumor epithelial cell proliferation, survival, and migration/invasion(52–54). It is also possible that PGE_2_ modulates disease through alteration of the microbiome, as NSAIDs have been implicated as potentially disrupting the gut microbiome(20, 55). Additionally, PGE_2_ functions as a key inflammatory signal that can regulate certain immune responses, with its local levels being tightly regulated during the trigger but also the resolution of inflammatory processes(56–58). Some of the best-known functions of PGE_2_ are indeed its role in intestinal inflammation and cancer, as well as its impact on the immune system(18). Paradoxically, we observed that pre-treatment with the COX inhibitor indomethacin caused a dysregulation of PG metabolism that led to increased PGE_2_ production upon CDI. This heightened PGE_2_ response was associated with elevated intestinal inflammatory cytokines, monocyte and neutrophil recruitment, partial dismantling of the intestinal epithelial cells tight junctions, and a specific disturbed microbiota composition.

Immune protection against *C. difficile* challenge seems to be independent of CD4^+^ cells, anti-toxin IgG and pIgR(59), but it strongly relies on rapid and effective myeloid cell responses (9, 37, 60). Cells of the immune system can exert critical roles in controlling bacterial pathogen damage and intestinal health through production of damaging or protective cytokines by T cells or Innate Lymphoid Cells (ILCs)(61, 62). Production of IFNγ by T cells and neutrophils has a protective role against CDI(63, 64), and the associated production of IL-12 by innate cells upon CDI can have a strong positive feedback effect on IFNγ production in this context. The role of IL17 cytokines and their cellular sources is more controversial, as they can induce damage but also trigger intestinal repair processes and maintain barrier integrity(65, 66). Perturbation of the microbiota induced by antibiotic treatment can also cause an imbalance of protective Treg:Th17 ratios(67). ILCs are however critical for controlling the acute response induced by CDI. In contrast to Rag1^-/-^ mice, Rag2^-/-^ Il2rγ^-/-^ mice rapidly succumb to CDI. While ILC3s display a limited role to resistance, loss of IFNγ expressing ILC1s in Rag1^-/-^ mice increased susceptibility(7). The contribution to CDI pathogenesis by other highly relevant cytokines like IL-23 and IL-22 provided by innate immune cells strongly depends on context(23, 24, 37, 68–70). In our studies, we found that indomethacin pre-treatment prior to CDI increased local levels of chemokines that induce recruitment of inflammatory myeloid cells like CXC2, CCL3 and CCL4, with a concomitant increase in circulating and local neutrophils numbers while type 3 ILC were unaltered. Also, in coordination with the increased levels of intestinal IL-6 and IL-1 β, total CD4^+^ cells and CD4^+^RORγt^+^ cells were found in larger numbers in the colonic lamina propria but not the draining mesenteric lymph nodes.

Intestinal epithelial cells constitute the main barrier against infectious agents colonizing the gastrointestinal tract. Cell junctional complexes, notably the tight junctions, regulate paracellular permeability and restrict the translocation of luminal microbes and microbial products across the epithelial monolayer(71). Displacement of occludin, but not ZO-1, from the junctions of mouse colonic epithelial cells during CDI did not manifest as gross morphological changes of TJ regions during EM visualization. This is reminiscent of the alterations seen in anti-CD3-, and TNF-α-treated mice, respectively, and consistent with the view that occludin is a regulator, rather than a key structural component, of TJs(72). However, indomethacin pretreatment with CDI redistributed both occludin and ZO-1 to the cytosol, and electron micrographs revealed a concomitant loss of TJ interactions. These changes are expected to increase paracellular permeability and promote bacterial translocation and could explain the observed increase in bacterial burden in the livers of indomethacin-treated, *C. difficile*-infected animals.

Induction of severe colitis upon CDI is subordinated to alterations in their microbiota caused by antibiotic administration that lead to dismantling of colonization resistance(5, 33, 37, 73, 74). Interestingly, recent studies have begun to highlight previously underappreciated and potentially detrimental effects of pharmaceutical drugs, such as NSAIDs, on the gut microbiota(20, 35). We observed that indomethacin did not cause an alteration of the microbial α-diversity but did induce significant alterations in microbiota structure that lead to an enrichment of *Bacteroides, Akkermansia, Porphyromonadaceae*, and *Parasutterella*, and a decrease in *Turicibacter* and *Porphyromonadaceae*. Interestingly, increases in *Bacteroides* and *Akkermansia* have been reported in association with inflammatory bowel disease and other infections(75–78). Furthermore, *Turicibacter* has been shown in several studies to be associated with colonization resistance to *C. difficile*(6, 79). Thus, indomethacin-mediated alterations in the microbiota may have a profound impact on manifestation and severity of CDI. Interestingly, when the microbiota α-diversity was severely reduced by antibiotic treatment, we found that *Paenibacillus* and *Akkermansia* expanded in mice pretreated with indomethacin. At present, the role of *Paenibacillus* in the pathology of CDI is unknown and further investigation is warranted.

Injury to intestinal epithelial barriers and microbial translocation can lead to a systemic response that mimics some aspects of sepsis and unveils a massive release of inflammatory cytokines, like IL-1β, and increases neutrophil and macrophage recruitment and activation (24). It is still unclear how the innate and adaptive arms of the immune response coordinate during CDI, especially in situations like the one we present with pre-treatment with indomethacin. In such circumstances, it is important to note that Type 3 ILCs can dysregulate adaptive immune CD4^+^ cell responses against commensal bacteria, but this ILC-mediated regulation of adaptive immune cells occurred independently of interleukin (IL)-17A, IL-22 or IL-23, but is dependent on antigen presentation(80). Additionally, antibiotics alone can promote inflammation through goblet cell-mediated translocation of native colonic microbiota in mice(81), but CDI induces significant goblet cell loss(82), which would counter effect this bacterial translocation in the experimental conditions such as those described in this work.

Our results highlight the capacity of a short-term oral dose of the NSAID indomethacin to cause an imbalance in PG production and disrupt the intestinal barrier to allow for bacterial entrance in the bloodstream. These effects are paralleled by a specific disarrangement of the intestinal microbiota and dysregulated inflammatory and immune responses that lead to increase pathological damage and finally unfold in an increased mortality rate. Our results call for caution in the use of NSAIDs in the context of *C. difficile* infections, but also potentially when other intestinal pathogens or insults co-occur with acute inflammatory events that affect PG balances. Moreover, we also highlight how a temporary modification of a set of key inflammatory mediators like PGs in the host can lead to significant perturbations to the resident gut microbiota. We believe that this unique combination of effects caused by indomethacin and CDI in the host and their microbiota could represent a generalized mechanism that leads to increased intestinal damage and complications when NSAIDs, or other drugs that alter key inflammatory molecules with pleiotropic effects, are used.

## Materials and Methods

### Experimental animals and infection model

All mice used in this study were obtained from Jackson laboratories (C57 BL6J) and were females of 6 weeks of age at arrival. Mice were given 2 weeks’ time to adapt to the new facilities and avoid stress associated responses and allow for in-house conditions adaptations. Mice were given Cefoperazone at 0.5 mg/ml in drinking water ad libitum for 5 days prior to treatment with Indomethacin (Cayman Chemical) at 10 mg/kg or vehicle (PBS) for 2 consecutive days by oral gave and then infected or not with 1×10^4^ spores of Clostridium difficile (NAP1/BI/027 strain M7404, O’Connor *et al.*, 2009) resuspended in PBS by oral gavage. Non-infected mice received only cefoperazone but afterwards only vehicle by oral gavage at the same time points. For some experiments untreated mice were used to obtain unaltered cecal microbiota.

### mRNA isolation, expression analysis and qRT-PCR

After bulk RNA isolation was isolated from tissues with Trizol (Life Technologies) following manufacturer’s instructions, a QIAgen RNeasy Plus Mini Kit was used to further purify mRNA for downstream analysis. mRNA expression was evaluated either using an nSolver inflammation panel mouse v2 from Nanostring directly on mRNA samples or by qRT-PCR performed using Applied Biosystems TaqMan amplification system after cDNA generation using a SuperScript VILO cDNA synthesis Kit (InVitrogen). qRT-PCR reactions, data quantification and analysis were performed in an Applied Biosystems 7300 Real Time PCR system using the following Taqman Primers: Reg3g (Mm00441127_m1), Muc2 (Mm01276696_m1), Ptger2 (Mm004360516_m 1), Ptger4 (Mm004360513_m1), Ptgs1 (Mm00477214_m1), Ptgs2 (Mm00478374_m1), Ptges (Mm00452105_m1), Hpgd (Mm00515121_m1), and Gapdh (Mm99999915_g1). Data generated by Nanostring technologies for mRNA quantification was analyzed with the company’s proprietary software nSolver 4.0.

### Bacterial burden in mouse organs

Liver was collected from the mouse with sanitized instruments and immediately placed in 1 mL of PBS in a 12 well plate. After the tissue was minced with scissors, 20 mL of the supernatant was drawn off and serially diluted. Dilutions were plated on TCCFA plates under aerobic and anaerobic conditions. After 24 hours, the plates were collected, CFUs were calculated and normalized to the weight of the liver. Cecum was also collected using sanitized instruments, and contents were expelled by placing pressure on the organ with a scalpel. Contents were then collected and put into a 1.5 ml tube. Weight of the contents was recorded, PBS was added, vortexed, and the slurry was serially diluted and plated onto TCCFA plates (Anaerobe Systems). After 24 hours, CFUs were counted and normalized to the weight of each sample.

### Tissue protein quantification and multiplex

Total cecum protein was isolated from ceca pre-washed with ice-cold PBS, homogenized a tissue shredder (Tissuemiser) and then centrifuged for 3 min at 8,600 g. Supernatants of these preparations were submitted for Luminex analysis of the provided analytes using x-map technology via the MapgPix^®^ system, in combination with multiplex kits from Millipore Sigma. Total tissue protein content was quantified by DC assay. Data was analyzed with GraphPad Prism 6.0, and heat maps were generated using the Morpheus software from the Broad Institute (https://software.broadinstitute.org/morpheus/).

### Flow cytometry

Cell suspensions from the mesenteric lymph nodes, peritoneal lavage and colon lamina propria from euthanized mice were obtained from mice at the indicated time points. Cell suspensions were incubated with Fc-block for 15 minutes on ice and then surface stained with a cocktail containing monoclonal anti mouse antibodies containing anti-CD19 (1D3), CD8a (5H10-1), anti-CD49b (DX5), anti-CD11b (M1/70), anti-CD11c (N418) and anti-CD196 (x29-2I.17) from Biolegend as well as anti-CD4 (RM4-5) and anti-Ly6G (1A8) from BD. After 30 minutes incubation on ice, cells were washed, fixed and permeabilized using the FoxP3 Fix/Perm buffer kit from eBiosciences/ThermoFisher and intracellular staining for RORγt (clone Q31-378) was performed as a last step. Flow cytometry data was obtained with BD FACSDiva™ 7.0 software and .fcs 3.0 files analyzed with FlowJo™.

### Electron and Immunofluorescence microscopy

Colonic tissue samples were fixed and stored in Karnovsky’s solution (4% paraformaldehyde in PBS (pH 7.4) with 1% glutaraldehyde) for at least 24 hrs at 4°C. Samples were neutralized with 125 mM glycine in PBS, post-fixed in 1% osmium tetroxide, and sequentially dehydrated with 15%, 30%, 50%, 70%, 90% and 100% ethanol. Samples were then infiltrated with Spurr’s Resin (Electron Microscopy Sciences, Hatfield, PA). Ultra-thin sections were contrasted with 2% uranyl acetate, followed by Reynold’s lead citrate and visualized with an FEI Tecnai Spirit transmission electron microscope (FEI, Hillsboro, OR) equipped with AMT CCD camera and AMT Image Capture Engine V602 software (Advanced Microscopy Techniques, Woburn, MA). For immunofluorescence microscopy tissue samples were frozen in OCT embedding medium (Tissue-Tek, Sakura Finetek, CA) and stored at −80°C. OCT-mounted tissue samples were sectioned at 3 μM thickness and fixed in 4% paraformaldehyde in PBS (pH 7.4) for 20 minutes at room temperature. Samples were washed with PBS, permeabilized with 0.2% Triton X-100 in PBS, quenched with 50 mM NH_4_Cl in PBS and then blocked with 5% IgG-free bovine serum albumin (BSA) in PBS. Primary antibodies used were 1:50 dilution of rabbit anti-occludin and rabbit anti-ZO1 (Abcam, Cambridge, MA). Samples were incubated with primary antisera overnight at 4°C, and then washed three times with 1% IgG-free BSA in PBS. Secondary antibodies (Alexa Fluor 555-conjugated anti-rabbit IgG) were added at 8 μg/ml in 5% IgG-free BSA for 1 hour. Samples were washed with PBS, stained with 4,6-diamidino-2-phenylindole (DAPI) and mounted in ProLong Diamond Antifade reagent (Thermo Fisher Scientific, Waltham, MA). Images were captured using DeltaVision Elite Deconvolution Microscope (GE Healthcare, Pittsburgh, PA) equipped with Olympus 100X/1.40 oil objective and using immersion oil (n=1.516) and GE Healthcare Software Version 6.5.2. ImageJ 1.51j8 (National Institutes of Health, Bethesda, MA) was used to merge and pseudocolor images.

### Colon histology and pathology scoring

Colons from experimental mice were collected at day 3 post-infection, then flushed with cold PBS, open longitudinally and rolled to generate swiss rolls. This colon rolls were fixed for 5 days in 10% buffered Formalin Phosphate and then transferred to 70% ethanol for 7 days. After that, these Swiss rolls were used to generate paraffin blocks that were stained with H&E and scored for the degree of injuries as described in Theriot CM et al., Gut Microbes 2:6, 326-34. 2011.

### DNA extraction, 16S rRNA gene sequencing, and gut microbiota analyses

Fecal samples were collected fresh from individual mice at prior to (baseline) and following treatment with cefoperazone, indomethacin, or a combination of cefoperazone and indomethacin. In a subset of mice (N=6/group), fecal samples were collected for the time course of a post-treatment 11-day recovery period. Following collection, fecal samples were immediately put on ice and subsequently frozen for storage at −20 °C. Microbial genomic DNA was extracted using the 96-well PowerSoil DNA isolation kit (Qiagen). For each sample, the V4 region of the bacterial 16S rRNA gene was amplified and sequenced using the Illumina MiSeq Sequencing platform as described elsewhere (Kozich et al., 2013). Sequences were curated using the mothur software package (v1.40.3) as previously described (Zackular et al., 2016; Schloss et al., 2009; Kozich et al., 2013). Briefly, the workflow we used included generating contigs with paired-end reads, filtering low quality sequences, aligning the resulting sequences to the SILVA 16S rRNA sequence database, and removed any chimeric sequences flagged by UCHIME. Following curation, we obtained between 9 and 83,525 sequences per sample (median = 13,161.5), with a median length of 253 bp. To minimize the impact of uneven sampling, the number of sequences in each sample was rarefied to 4200. Sequences were clustered into OTUs based on a 3% distance cutoff calculated using the OptiClust algorithm. All sequences were classified using the Ribosomal Database Project training set (version 16) and OTUs were assigned a taxonomic classification using a naive Bayesian classifier. Significantly altered OTUs for each group were selected using the biomarker discovery algorithm LEfSe (linear discriminant analysis (LDA) effect size) in mothur (Segata et al., 2011). α-diversity was calculated using the Shannon diversity index and β-diversity was calculated using the θ_YC_ distance metric with OTU frequency data. FASTQ sequence data obtained in this study has been deposited to the Sequence Read Archive (SRA) at NCBI under the accession number SRP152292.

## Authors contributions

D.M.A., D.M., and J.P.Z. designed the research and analyzed the data. D.M., J.P.Z., B.T., L.K., J.L.R., L.M.R., and M.K.W. performed the experiments, J.P.Z. generated the microbiota libraries and performed the corresponding bioinformatics analysis. V.K.V., G.V., L.J.C. and P.S., helped in with data analysis and interpretation. D.M., J.P.Z., and D.M.A. wrote the manuscript, and J.L.R., V.K.V., and E.S. proofread the manuscript.

## References

1. Lessa FC, Gould C V., Clifford McDonald L. 2012. Current status of clostridium difficile infection epidemiology. Clin Infect Dis.

2. Leffler DA, Lamont JT. 2015. Clostridium difficile infection. N Engl J Med 372:1539–48.

3. Buonomo EL, Petri WA. 2016. The microbiota and immune response during Clostridium difficile infection. Anaerobe 41:79–84.

4. Lessa FC, Mu Y, Bamberg WM, Beldavs ZG, Dumyati GK, Dunn JR, Farley MM, Holzbauer SM, Meek JI, Phipps EC, Wilson LE, Winston LG, Cohen JA, Limbago BM, Fridkin SK, Gerding DN, McDonald LC. 2015. Burden of *Clostridium difficile* Infection in the United States. N Engl J Med 372:825–834.

5. Collins J, Robinson C, Danhof H, Knetsch CW, Van Leeuwen HC, Lawley TD, Auchtung JM, Britton RA. 2018. Dietary trehalose enhances virulence of epidemic Clostridium difficile. Nature 553:291–294.

6. Zackular JP, Moore JL, Jordan AT, Juttukonda LJ, Noto MJ, Nicholson MR, Crews JD, Semler MW, Zhang Y, Ware LB, Washington MK, Chazin WJ, Caprioli RM, Skaar EP. 2016. Dietary zinc alters the microbiota and decreases resistance to Clostridium difficile infection. Nat Med 22:1330–1334.

7. Abt MC, Lewis BB, Caballero S, Xiong H, Carter RA, Susac B, Ling L, Leiner I, Pamer EG. 2015. Innate immune defenses mediated by two ilc subsets are critical for protection against acute clostridium difficile infection. Cell Host Microbe 18:27–37.

8. Garey KW, Jiang Z-D, Ghantoji S, Tam VH, Arora V, Dupont HL. 2010. A common polymorphism in the interleukin-8 gene promoter is associated with an increased risk for recurrent Clostridium difficile infection. Clin Infect Dis 51:1406–10.

9. Jarchum I, Liu M, Shi C, Equinda M, Pamer EG. 2012. Critical role for myd88-Mediated Neutrophil recruitment during Clostridium difficile colitis. Infect Immun 80:2989–2996.

10. Madan R, Petri WA. 2012. Immune responses to Clostridium difficile infection. Trends Mol Med 18:658–666.

11. Permpalung N, Upala S, Sanguankeo A, Sornprom S. 2016. Association between NSAIDs and Clostridium difficile-Associated Diarrhea: A Systematic Review and Meta-Analysis. Can J Gastroenterol Hepatol 2016:7431838.

12. Liang X, Bittinger K, Xuanwen L, Abernethy DR, Bushman FD, Fitzgerald GA. 2015. Bidirectional interactions between indomethacin and the murine intestinal microbiota. Elife 4.

13. Xiao X, Nakatsu G, Jin Y, Wong S, Yu J, Lau JYW. 2017. Gut Microbiota Mediates Protection Against Enteropathy Induced by Indomethacin. Sci Rep 7.

14. Cryer B, Barnett MA, Wagner J, Wilcox CM. 2016. Overuse and Misperceptions of Nonsteroidal Anti-inflammatory Drugs in the United States. Am J Med Sci 352:472–480.

15. Fowler TO, Durham CO, Planton J, Edlund BJ. 2014. Use of nonsteroidal anti-inflammatory drugs in the older adult. J Am Assoc Nurse Pract 26:414–423.

16. Tonolini M. 2013. Acute nonsteroidal anti-inflammatory drug-induced colitis. J Emerg Trauma Shock 6:301–3.

17. Villanacci V, Casella G, Bassotti G. 2011. The spectrum of drug-related colitides: Important entities, though frequently overlooked. Dig Liver Dis 43:523–528.

18. Montrose DC, Nakanishi M, Murphy RC, Zarini S, McAleer JP, Vella AT, Rosenberg DW. 2015. The role of PGE2 in intestinal inflammation and tumorigenesis. Prostaglandins Other Lipid Mediat 116–117:26–36.

19. Blackler RW, De Palma G, Manko A, Da Silva GJ, Flannigan KL, Bercik P, Surette MG, Buret AG, Wallace JL. 2015. Deciphering the pathogenesis of NSAID enteropathy using proton pump inhibitors and a hydrogen sulfide-releasing NSAID. Am J Physiol Liver Physiol 308:G994–G1003.

20. Rogers MAM, Aronoff DM. 2015. The Influence of Nonsteroidal Anti-Inflammatory Drugs on the Gut Microbiome. Clin Microbiol Infect 22:178.e1–178.e9.

21. Syer SD, Blackler RW, Martin R, de Palma G, Rossi L, Verdu E, Bercik P, Surette MG, Aucouturier A, Langella P, Wallace JL. 2015. NSAID enteropathy and bacteria: a complicated relationship. J Gastroenterol.

22. X. X, D. N, Y. J, J. Y, J.Y. L. 2015. Beneficial changes in intestinal microbiota during indomethacin treatment in mice. Gastroenterology 148:S730.

23. Buonomo EL, Madan R, Pramoonjago P, Li L, Okusa MD, Petri WA. 2013. Role of interleukin 23 signaling in clostridium difficile colitis, p. 917–920. In Journal of Infectious Diseases.

24. Hasegawa M, Kamada N, Jiao Y, Liu MZ, Núñez G, Inohara N. 2012. Protective role of commensals against Clostridium difficile infection via an IL-1β-mediated positive-feedback loop. J Immunol 189:3085–91.

25. Jafari N V., Kuehne SA, Bryant CE, Elawad M, Wren BW, Minton NP, Allan E, Bajaj-Elliott M. 2013. Clostridium difficile Modulates Host Innate Immunity via Toxin-Independent and Dependent Mechanism(s). PLoS One 8.

26. Kamada N, Chen GY, Inohara N, Núñez G. 2013. Control of pathogens and pathobionts by the gut microbiota. Nat Immunol 14:685–690.

27. Bjarnason I, Zanelli G, Smith T, Prouse P, Williams P, Smethurst P, Delacey G, Gumpel MJ, Levi AJ. 1987. Nonsteroidal antiinflammatory drug-induced intestinal inflammation in humans. Gastroenterology 93:480–9.

28. Berg DJ, Zhang J, Weinstock J V., Ismail HF, Earle KA, Alila H, Pamukcu R, Moore S, Lynch RG. 2002. Rapid development of colitis in NSAID-treated IL-10-deficient mice. Gastroenterology 123:1527–1542.

29. Feuerstadt P. 2015. Clostridium difficile Infection. Clin Transl Gastroenterol 6.

30. Mukherjee S, Hooper L V. 2015. Antimicrobial Defense of the Intestine. Immunity 42:28–39.

31. Rao K, Erb-Downward JR, Walk ST, Micic D, Falkowski N, Santhosh K, Mogle JA, Ring C, Young VB, Huffnagle GB, Aronoff DM. 2014. The systemic inflammatory response to clostridium difficile infection. PLoS One 9.

32. Shi C, Pamer EG. 2014. Monocyte Recruitment Suring Infection and Inflammation. Nat Rev Immunol 11:762–774.

33. Buffie CG, Jarchum I, Equinda M, Lipuma L, Gobourne A, Viale A, Ubeda C, Xavier J, Pamer EG. 2012. Profound alterations of intestinal microbiota following a single dose of clindamycin results in sustained susceptibility to Clostridium difficile-induced colitis. Infect Immun 80:62–73.

34. Jenior ML, Leslie JL, Young VB, Schloss PD. 2017. *Clostridium difficile* Colonizes Alternative Nutrient Niches during Infection across Distinct Murine Gut Microbiomes. mSystems 2:e00063–17.

35. Maier L, Pruteanu M, Kuhn M, Zeller G, Telzerow A, Anderson EE, Brochado AR, Fernandez KC, Dose H, Mori H, Patil KR, Bork P, Typas A. 2018. Extensive impact of non-antibiotic drugs on human gut bacteria. Nature 555:623–628.

36. Evans CT, Safdar N. 2015. Current trends in the epidemiology and outcomes of clostridium difficile infection. Clin Infect Dis 60:S66–S71.

37. Abt MC, McKenney PT, Pamer EG. 2016. Clostridium difficile colitis: Pathogenesis and host defence. Nat Rev Microbiol.

38. Olsen MA, Yan Y, Reske KA, Zilberberg MD, Dubberke ER. 2015. Recurrent Clostridium difficile infection is associated with increased mortality. Clin Microbiol Infect 21:164–170.

39. Newman KM, Rank KM, Vaughn BP, Khoruts A. 2017. Treatment of recurrent Clostridium difficile infection using fecal microbiota transplantation in patients with inflammatory bowel disease. Gut Microbes.

40. Smits LP, Bouter KEC, De Vos WM, Borody TJ, Nieuwdorp M. 2013. Therapeutic potential of fecal microbiota transplantation. Gastroenterology 145:946–953.

41. Bojanova DP, Bordenstein SR. 2016. Fecal Transplants: What Is Being Transferred? PLoS Biol 14.

42. Zhou Y, Boudreau DM, Freedman AN. 2014. Trends in the use of aspirin and nonsteroidal anti-inflammatory drugs in the general U.S. population. Pharmacoepidemiol Drug Saf 23:43–50.

43. Ananthakrishnan AN, Higuchi LM, Huang ES, Khalili H, Richter JM, Fuchs CS, Chan AT. 2012. Aspirin, nonsteroidal anti-inflammatory drug use, and risk for Crohn disease and ulcerative colitis: a cohort study. Ann Intern Med 156:350–359.

44. Kelsen JR, Kim J, Latta D, Smathers S, McGowan KL, Zaoutis T, Mamula P, Baldassano RN. 2011. Recurrence rate of clostridium difficile infection in hospitalized pediatric patients with inflammatory bowel disease. Inflamm Bowel Dis 17:50–55.

45. Kvasnovsky CL, Aujla U, Bjarnason I. 2014. Nonsteroidal anti-inflammatory drugs and exacerbations of inflammatory bowel disease. Scand J Gastroenterol 50:255–263.

46. Sokol H, Lalande V, Landman C, Bourrier A, Nion-Larmurier I, Rajca S, Kirchgesner J, Seksik P, Cosnes J, Barbut F, Beaugerie L. 2017. Clostridium difficile infection in acute flares of inflammatory bowel disease: A prospective study. Dig Liver Dis 49:643–646.

47. Kim H, Rhee SH, Pothoulakis C, LaMont JT. 2007. Inflammation and Apoptosis in Clostridium difficile Enteritis Is Mediated by PGE2 Up-Regulation of Fas Ligand. Gastroenterology 133:875–886.

48. Miyoshi H, VanDussen KL, Malvin NP, Ryu SH, Wang Y, Sonnek NM, Lai C, Stappenbeck TS. 2017. Prostaglandin E2 promotes intestinal repair through an adaptive cellular response of the epithelium. EMBO J 36:5–24.

49. Roulis M, Nikolaou C, Kotsaki E, Kaffe E, Karagianni N, Koliaraki V, Salpea K, Ragoussis J, Aidinis V, Martini E, Becker C, Herschman HR, Vetrano S, Danese S, Kollias G. 2014. Intestinal myofibroblast-specific Tpl2-Cox-2-PGE2 pathway links innate sensing to epithelial homeostasis. Proc Natl Acad Sci 111:E4658–E4667.

50. Stenson WF. 2007. Prostaglandins and epithelial response to injury. Curr Opin Gastroenterol.

51. Wells JM, Rossi O, Meijerink M, van Baarlen P. 2011. Epithelial crosstalk at the microbiota-mucosal interface. Proc Natl Acad Sci 108:4607–4614.

52. Lanas A, Sopeña F. 2009. Nonsteroidal Anti-Inflammatory Drugs and Lower Gastrointestinal Complications. Gastroenterol Clin North Am.

53. Nakanishi M, Rosenberg DW. 2013. Multifaceted roles of PGE2 in inflammation and cancer. Semin Immunopathol.

54. Schumacher Y, Aparicio T, Ourabah S, Baraille F, Martin A, Wind P, Dentin R, Postic C, Guilmeau S. 2016. Dysregulated CRTC1 activity is a novel component of PGE2 signaling that contributes to colon cancer growth. Oncogene 35:2602–2614.

55. van Opstal E, Kolling GL, Moore JH, Coquery CM, Wade NS, Loo WM, Bolick DT, Shin JH, Erickson LD, Warren CA. 2016. Vancomycin Treatment Alters Humoral Immunity and Intestinal Microbiota in an Aged Mouse Model of *Clostridium difficile* Infection. J Infect Dis 214:jiw071.

56. Agard M, Asakrah S, Morici LA. 2013. PGE(2) suppression of innate immunity during mucosal bacterial infection. Front Cell Infect Microbiol 3:45.

57. Duffin R, OConnor RA, Crittenden S, Forster T, Yu C, Zheng X, Smyth D, Robb CT, Rossi F, Skouras C, Tang S, Richards J, Pellicoro A, Weller RB, Breyer RM, Mole DJ, Iredale JP, Anderton SM, Narumiya S, Maizels RM, Ghazal P, Howie SE, Rossi AG, Yao C. 2016. Prostaglandin E2 constrains systemic inflammation through an innate lymphoid cell-IL-22 axis. Science (80-) 351:1333–1338.

58. Yao C, Sakata D, Esaki Y, Li Y, Matsuoka T, Kuroiwa K, Sugimoto Y, Narumiya S. 2009. Prostaglandin E2–EP4 signaling promotes immune inflammation through TH1 cell differentiation and TH17 cell expansion. Nat Med 15:633–640.

59. Johnston PF, Gerding DN, Knight KL. 2014. Protection from clostridium difficile infection in CD4 T cell- and polymeric immunoglobulin receptor-deficient mice. Infect Immun 82:522–531.

60. Duque GA, Descoteaux A. 2014. Macrophage cytokines: Involvement in immunity and infectious diseases. Front Immunol.

61. Mcsorley SJ. 2014. Immunity to intestinal pathogens: Lessons learned from Salmonella. Immunol Rev.

62. Sonnenberg GF, Artis D. 2015. Innate lymphoid cells in the initiation, regulation and resolution of inflammation. Nat Med.

63. Ishida Y, Maegawa T, Kondo T, Kimura A, Iwakura Y, Nakamura S, Mukaida N. 2004. Essential involvement of IFN-gamma in Clostridium difficile toxin A-induced enteritis. J Immunol 172:3018–25.

64. Yu H, Chen K, Sun Y, Carter M, Garey KW, Savidge TC, Devaraj S, Tessier ME, Von Rosenvinge EC, Kelly CP, Pasetti MF, Feng H. 2017. Cytokines are markers of the Clostridium difficile-induced inflammatory response and predict disease severity. Clin Vaccine Immunol 24.

65. Lee JS, Tato CM, Joyce-Shaikh B, Gulan F, Cayatte C, Chen Y, Blumenschein WM, Judo M, Ayanoglu G, McClanahan TK, Li X, Cua DJ. 2015. Interleukin-23-Independent IL-17 Production Regulates Intestinal Epithelial Permeability. Immunity 43:727–738.

66. McDermott AJ, Falkowski NR, McDonald RA, Pandit CR, Young VB, Huffnagle GB. 2016. Interleukin-23 (IL-23), independent of IL-17 and IL-22, drives neutrophil recruitment and innate inflammation during Clostridium difficile colitis in mice. Immunology 147:114–124.

67. Becattini S, Taur Y, Pamer EG. 2016. Antibiotic-Induced Changes in the Intestinal Microbiota and Disease. Trends Mol Med xx:1–21.

68. Aden K, Rehman A, Falk-Paulsen M, Secher T, Kuiper J, Tran F, Pfeuffer S, Sheibani-Tezerji R, Breuer A, Luzius A, Jentzsch M, Häsler R, Billmann-Born S, Will O, Lipinski S, Bharti R, Adolph T, Iovanna JL, Kempster SL, Blumberg RS, Schreiber S, Becher B, Chamaillard M, Kaser A, Rosenstiel P. 2016. Epithelial IL-23R Signaling Licenses Protective IL-22 Responses in Intestinal Inflammation. Cell Rep 16:2208–2218.

69. Cowardin CA, Kuehne SA, Buonomo EL, Marie CS, Minton NP, Petri WA. 2015. Inflammasome activation contributes to interleukin-23 production in response to Clostridium difficile. MBio 6.

70. Lee YS, Yang H, Yang JY, Kim Y, Lee SH, Kim JH, Jang YJ, Vallance BA, Kweon MN. 2015. Interleukin-1 (IL-1) signaling in intestinal stromal cells controls KC/ CXCL1 secretion, which correlates with recruitment of IL-22- secreting neutrophils at early stages of Citrobacter rodentium infection. Infect Immun 83:3257–3267.

71. Roxas JL, Viswanathan VK. 2018. Modulation of intestinal paracellular transport by bacterial pathogens. Compr Physiol 8:823–842.

72. Marchiando AM, Shen L, Vallen Graham W, Weber CR, Schwarz BT, Austin JR, Raleigh DR, Guan Y, Watson AJM, Montrose MH, Turner JR. 2010. Caveolin-1-dependent occludin endocytosis is required for TNF-induced tight junction regulation in vivo. J Cell Biol 189:111–126.

73. Chen X, Katchar K, Goldsmith JD, Nanthakumar N, Cheknis A, Gerding DN, Kelly CP. 2008. A mouse model of Clostridium difficile-associated disease. Gastroenterology 135:1984–1992.

74. Theriot CM, Koumpouras CC, Carlson PE, Bergin II, Aronoff DM, Young VB. 2011. Cefoperazone-treated mice as an experimental platform to assess differential virulence of Clostridium difficile strains. Gut Microbes 2:326–334.

75. Imhann F, Vila AV, Bonder MJ, Fu J, Gevers D, Visschedijk MC, Spekhorst LM, Alberts R, Franke L, van Dullemen HM, Ter Steege RWF, Huttenhower C, Dijkstra G, Xavier RJ, Festen EAM, Wijmenga C, Zhernakova A, Weersma RK. 2016. Interplay of host genetics and gut microbiota underlying the onset and clinical presentation of inflammatory bowel disease. Gut.

76. Pascal V, Pozuelo M, Borruel N, Casellas F, Campos D, Santiago A, Martinez X, Varela E, Sarrabayrouse G, Machiels K, Vermeire S, Sokol H, Guarner F, Manichanh C. 2017. A microbial signature for Crohn’s disease. Gut 66:813–822.

77. Seregin SS, Golovchenko N, Schaf B, Chen J, Pudlo NA, Mitchell J, Baxter NT, Zhao L, Schloss PD, Martens EC, Eaton KA, Chen GY. 2017. NLRP6 Protects II10−/−Mice from Colitis by Limiting Colonization of Akkermansia muciniphila. Cell Rep 19:733–745.

78. Tedjo DI, Smolinska A, Savelkoul PH, Masclee AA, Van Schooten FJ, Pierik MJ, Penders J, Jonkers DMAE. 2016. The fecal microbiota as a biomarker for disease activity in Crohn’s disease. Sci Rep 6.

79. Schubert AM, Rogers M a M, Ring C, Mogle J, Petrosino JP, Young VB, Aronoff DM, Schloss PD. 2014. Microbiome Data Distinguish Patients with Clostridium difficile Infection and Non- C. difficile-Associated Diarrhea from Healthy. MBio 5:1–9.

80. Hepworth MR, Monticelli LA, Fung TC, Ziegler CGK, Grunberg S, Sinha R, Mantegazza AR, Ma H-L, Crawford A, Angelosanto JM, Wherry EJ, Koni PA, Bushman FD, Elson CO, Eberl G, Artis D, Sonnenberg GF. 2013. Innate lymphoid cells regulate CD4+ T-cell responses to intestinal commensal bacteria. Nature 498:113–117.

81. Knoop K a., McDonald KG, Kulkarni DH, Newberry RD. 2015. Antibiotics promote inflammation through the translocation of native commensal colonic bacteria. Gut 1–10.

82. Batah J, Kobeissy H, Pham PTB, Denève-Larrazet C, Kuehne S, Collignon A, Janoir-Jouveshomme C, Marvaud JC, Kansau I. 2017. Clostridium difficile flagella induce a pro-inflammatory response in intestinal epithelium of mice in cooperation with toxins. Sci Rep 7.

